# The N501Y and K417N mutations in the spike protein of SARS-CoV-2 alter the interactions with both hACE2 and human derived antibody: A Free energy of perturbation study

**DOI:** 10.1101/2020.12.23.424283

**Authors:** Filip Fratev

## Abstract

The N501Y and K417N mutations in spike protein of SARS-CoV-2 and their combination arise questions but the data about their mechanism of action at molecular level is limited. Here, we present Free energy perturbation (FEP) calculations for the interactions of the spike S1 receptor binding domain (RBD) with both the ACE2 receptor and an antibody derived from COVID-19 patients. Our results shown that the S1 RBD-ACE2 interactions were significantly increased whereas those with the STE90-C11 antibody dramatically decreased; about over 100 times. The K417N mutation had much more pronounced effect and in a combination with N501Y fully abolished the antibody effect. This may explain the observed in UK and South Africa more spread of the virus but also raise an important question about the possible human immune response and the success of already available vaccines.

## Letter

A discussion about the N501Y and K412N mutations in COVID-19 continue to arise many question but little data are currently available [1–2]. During the last weeks, the N501Y mutation (B.1.1.7 lineage) has been mainly observed in UK whereas the combination of N501Y and K417N mutations (501Y.V2 lineage) in a South Africa (SA). This led to new restrictions and many countries closed their borders for the travelers coming from the island. A little is known about the N501Y and K417N but their positions and well established interaction with the human ACE2 protein (hACE2), which is responsible for the virus entry into cells, deserve a special attention. Moreover, it has been shown that N501Y significantly increases virus adaption in a mouse model [3]. In addition, an enhancement of the virus transmission in humans of about 70% and even worse were reported [1–2, 4–5]. Thus, data about the molecular mechanism of action N501Y and K417N are urgently need.

Despite the computational demands, the Free energy of binding (FEP) approach is one of the most successful and precise *in silico* techniques for accurate prediction of both the ligand selectivity [6–7], protein-protein interactions [8–9] and protein stability [10–11]. It outperforms significantly the traditional molecular dynamics based methodologies, such as for example MM/GBSA and empirical solutions like FoldX and etc. and often precisely predicts the free energy differences between the mutations with a RMSE of about only1.2 kcal/mol [10–11]. It has been also shown that the better sampling approaches lead to much better results and most importantly for more than 90% correct predictions of the effect; i.e. whether the effect will be positive or negative after certain ligand or protein substitutions [12].

To calculate the differences in the free energy of binding for each complex in this study we employed the Desmond FEP/REST approach described in details previously [5–12]. Initially, the default sampling protocol was applied with the number of lambda (λ) windows either 12 or 24, in dependence of the mutation charge. An equilibration and 5 ns-long replica exchange solute tempering (REST) simulations in a muVT ensemble was further conducted. Only the mutated atoms was included in the REST “hot region” region. OPLS3e force field was used for the all simulations [13]. A set of N501 mutations for which experimental data is available was selected to validate the calculations. The experimental structure of the S1 RBD-hACE2 complex (pdb: 6M0J) was used as a starting point. After the solvation with SPC waters the complex consisted of over 102 000 atoms. For the study of S1 RBD interactions with the neutralizing antibody STE90-C11 selected from COVID-19 patients [14], which was well tolerated to the known mutations, we use the very recently published X-ray structure with a PDB access number of 7B3O. For the FEP calculations with double N501Y/K417N mutations we used as a starting point the energy minimized most representative frame of the N501Y FEP simulation.

**Table 1** presents the results from FEP calculations. As one can see the energy convergences in some cases were not so good for both the Bennett and the cycle closure (CC) approaches (see **Figures 1A**). In fact, the error of the calculated ΔΔGCC predictions, during the first 5 ns, suffered from much larger standard deviations. The convergence is indeed a very important issue during the FEP calculations. Hence, we are extended the FEP calculations to 20 ns-long REST sampling and obtained a good convergence (see **Table 1** and **Figure 1D**). Another set of simulations are currently undergoing in our lab performed by our own developed sampling protocol [12]. In the particular case, before the REST procedure we use an equilibration of each λ for 50 ns and then run 20 ns-long REST simulations.

**Table 1.** FEP results for the selected mutations. See the text for details. CCC is the cycle closure (CC) convergence

| Mutation | $\Delta\Delta G$<br>Bennett<br>(5 ns) | Energy<br>Conv.<br>5 ns | CCC<br>5 ns | $\Delta\Delta G$<br>Bennett<br>(15 ns) | $\Delta\Delta G$ CC<br>(15 ns) | Energy<br>Conv.<br>15 ns | CCC<br>15 ns |
| --- | --- | --- | --- | --- | --- | --- | --- |
| N501Y-STE90-C11 | 3.78 (0.15) | Good | N/A | 3.48 (0.12) | N/A | Good | N/A |
| K417N-STE90-C11 | 5.64 (0.17) | Good | N/A | 5.74 (0.15) | N/A | Good | N/A |
| N501Y/K417N-STE90-C11 | 8.61 (0.46) | Fair | N/A | 5.83 (0.43) | N/A | Good | N/A |
| K417N-ACE2 | -0.39 (0.23) | Fair | N/A | N/A | N/A | Good | N/A |
| N501Y-ACE2 | -0.5 (0.34) | Fair | Fair | -1.75 (0.37) | -1.42 (0.56) | Good | Good |
| N501D-ACE2 | 6.25 (0.46) | Bad | Bad | 3.54 (0.44) | 3.22 (0.56) | Good | Good |
| N501T-ACE2 | -1.69 (0.29) | Fair | Fair | -1.90 (0.17) | -2.24 (0.60) | Good | Good |
| N501F | 0.78 (0.27) | Bad | Bad | -0.00 (0.33) | 0.34 (0.60) | Good | Good |

**Figure 1.**
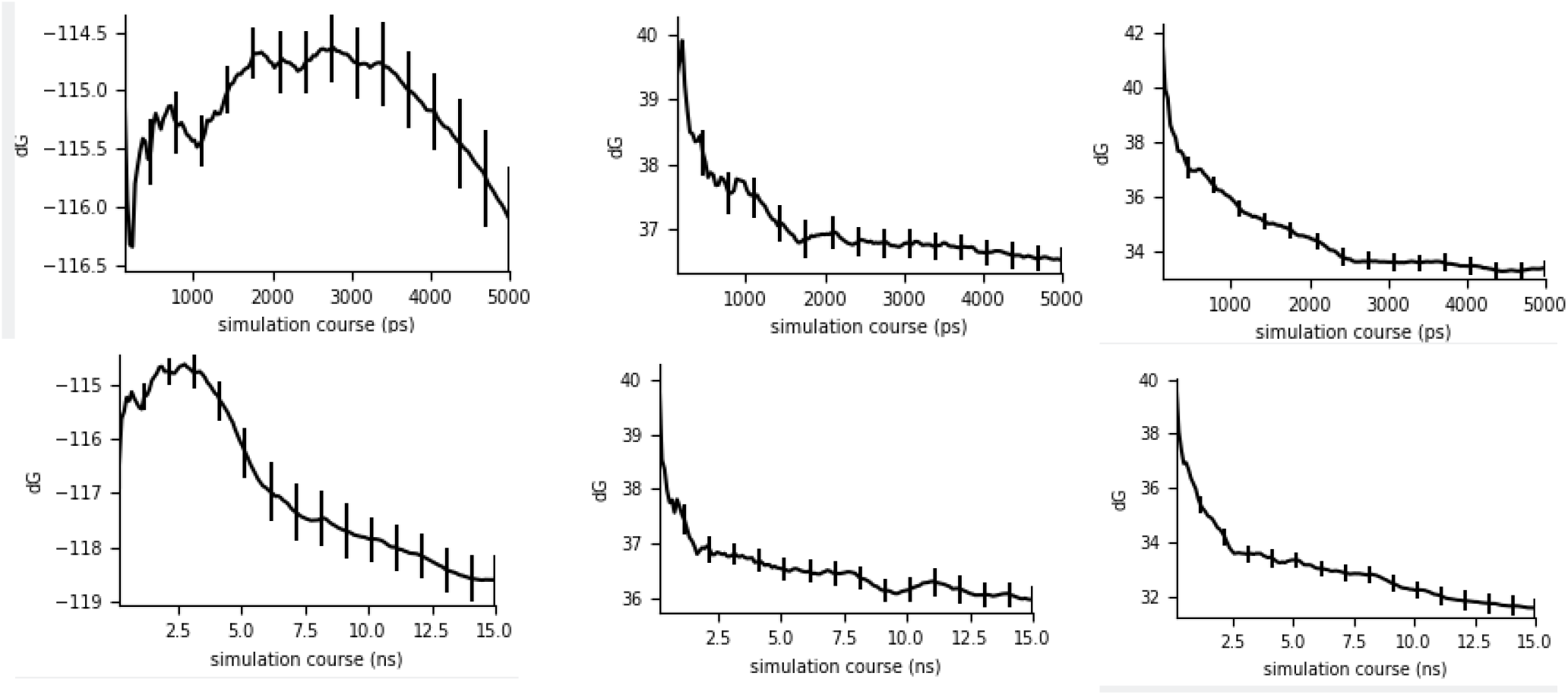
Observed convergence of the calculated Bennett free energies for the complex legs of the mutations: ***(A)*** N501D-ACE2, ***(B)*** N501Y-STE90-C11 and ***(C)*** N501Y-ACE2, after 5 ns FEP, respectively and (***D***) N501D-ACE2 (***E***) N501Y-STE90-C11 and (***F***) after 15ns of the FEP simulations.

The cycle closure calculations for protein-protein selectivity have been not studied well, thus, to avoid any confusions in the interpretations we used only the Bennett values to make our conclusions herein. Moreover, during the initial phase of our study the K417N mutation was not reported yet and it is a difficult to obtain the ΔΔGCC values for the S1 RBD-STE90-C11 complex at this stage. The better sampling protocols could make the free energies more precise and improve the CC ΔΔG values, however, it is not likely to change qualitatively the results present here; i.e. whether one mutation can results either in an increase or decrease in the interactions. This conclusion is also based on our experience with such type of simulations with both the ligand-protein complexes and protein mutations [12, 15–16].

The main results of the FEP study were:

1. We observed significant decrease of the binding between S1 RBD and STE90-C11 antibody by ΔΔG of 3.78 kcal/mol. This is a significant value and the observed convergence was good (**Table 1** and **Figures 1B** and **1E**). The binding energy of the antibody can be roughly estimated based on the published value of IC_50_=0.56nM in a plaque-based live SARS-CoV-2 neutralization assay [14]. This is equivalent to ΔG value of about −12.7 kcal/mol (ΔG=RTln (IC_50_). Thus, our calculations predict that the N501Y mutant will produce a decrease of the binding into ΔG = −8.92 kcal/mol or 293 nM. This is about 161 times lower than the wild type. The value from extended 15 ns-long FEP simulation was similar: **3.48 kcal/mol**.
2. Initially, we detected an increase of the binding between S1 RBD and human ACE2 by ΔΔG value of only −0.5 kcal/mol after 5 ns of FEP calculations. However, considering the decreasing free energy trend of the complex leg (see **Figure 1C**; it decreased by about 1 kcal/mol only for the last 2 ns of the FEP simulations) and 500 ns-long classical MD simulation we expected this value to be much more significant after extension of the REST sampling to 15 ns. This hypothesis was confirmed and we obtained a ΔΔG value of **−1.75 kcal/mol** from the extended simulation and it will eventually go down a bit more because the convergence is still not perfect (**Figure 1F**). In support to our data are also the *in vivo* studies of N501Y on mice [3].
3. The addition of K417N mutation led to a dramatic decrease of the STE90-C11 antibody binding to virus’s S1 RBD. The default FEP sampling protocol calculated a ΔΔG value of **5.74 kcal/mol**. In the case when both N501Y and K417N mutations are present at the same time this value further increased to 8.61 kcal/mol but this simulation was not well converged and after its extension to 15 ns we obtained a valued of **5.83 kcal/mol**. Thus, it seems that the effect of these mutations is not additive and only the K417N mutation can abolish the interaction with STE90-C11 antibody. These results also suggest that even the well tolerated to mutations antibodies eventually would be resisted to this variant of SARS-Cov-2.
4. The K417N mutation also increases the S1 RBD binding to ACE2 by **−0.39 kcal/mol**. The convergence after 5 ns of FEP/REST simulations was better than in the N501Y case but we expect a further decrease in the ΔΔG value after the 15 ns extension. Thus, both the N501Y and K417N mutations enhanced the RBD binding to ACE2.

These results can be considered as a trustful because reproduced well the experimental data [ref. 17, see Fig 4A]. For instance, for the N501D mutation we calculated a ΔΔG value of **3.54 kcal/mol** meaning that it greatly reduces the S1 RBD binding to ACE2, in accordance to the experimentally observed change of over 100 times. In contrary, the N501T mutation transformed the S1 RBD to a better binder (ΔΔG = **−1.69 kcal/mol**), in an excellent agreement with the experimental data. Additional sets of calculations for other mutations, such as for example the transformation of N501D to N501Y (ΔΔG = **−4.32 kcal/mol**), were also performed in order the CC ΔΔG predictions to be obtained. The latter residue substitution provides an additional support that the N501Y mutation increases significantly the binding. In conclusion, it is evident from both the experimental data and FEP study here that the binding of the spike S1 RBD to hACE2 is highly sensitive to the N501 mutations and even the substitutions with small residues, such as N501T, seems to alter the structure of the complex. Based on the all FEP calculations it is also evident that at least 15-20 ns –long FEP/REST simulations for protein selectivity are required for systems with more than 100 000 atoms.

Further, we identified the conformational changes in the N501Y mutated S1 RBD-ACE2 complex. In particular, we compared the most representative structures after 5 ns of FEP simulation for the wild type and the average structure obtained by 500 ns-long classical MD simulation (**Figure 2A**). As one can see, the mutated S1 RBD rotates by about 20 degree. In a result, the RBD can approach deeper into the center of the binding site with ACE2 and the distance between Cα atoms of Tyr501 and Lys352 decreased by about 1.0 Å. The position of the surface residues was also changed. The Tyr501 makes a stable H-bond with the crucial for the ACE2 binding residue Lys353 (**Figure 2B**) but the longer 500 ns-long MD simulation showed that this bond is not so pronounced and mainly the hydrophobic and π-π stacking of Tyr501 increase the binding strength. The Leu455 was in a much closer position to ACE2 interacting with the surface helix residues. A conformational change of other residues were also detected as such for example Thr500, Tyr505, Tyr449, Tyr453, Gln493 and other. These simulations explain also why the G502 and L455 mutations are so sensitive to the ACE2 interactions.

**Figure 2.**
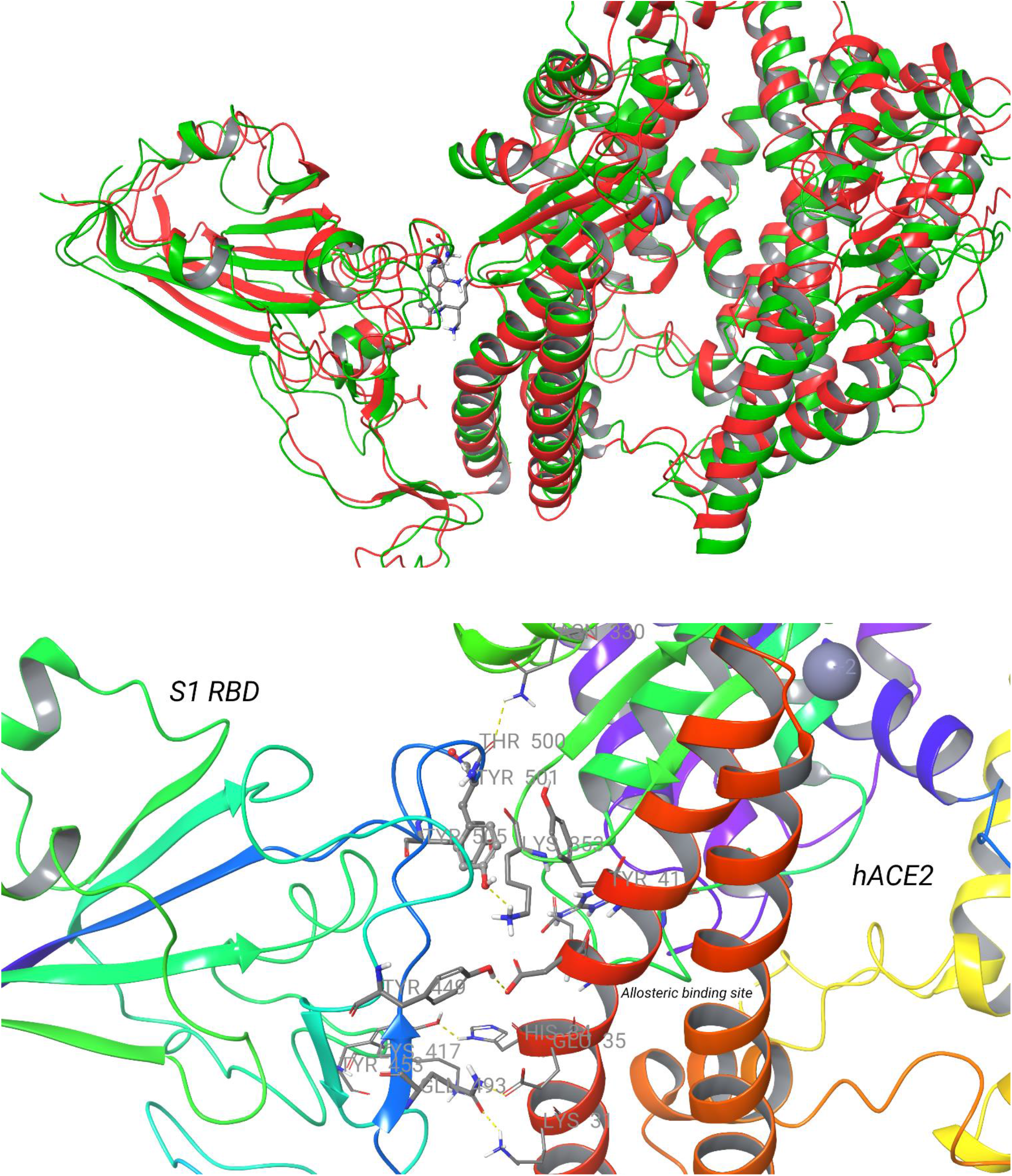
The identified changes in the interactions between S1 RDB and hACE2. ***(A)*** An alignment of the obtained structures after FEP 5ns-long simulation of WT (5ns; in green color) and 500 ns-long classical MD of N501Y mutant (in red color). ***(B)*** A close look of identified by FEP simulation interactions of N501Y mutant. With yellow dot lines are shown the H-bonds.

Recently we identified a set of ACE2 allosteric modulators which bind to ACE2 but did not produce any significant reduction in the virus replication (unpublished results). We targeted into the binding site located inside ACE2 which is close to the virus’s S1 RBD (see **Figure 2B**). Thus, it seems that the virus can act in a different way, as such for example entry mediation by Neuropilin-1 [18] or other processes are also possible. To our best knowledge there are no other similar ACE2 binders developed up to the moment which are able to reduce the virus replication, not only the interactions with S1 RBD. However, it is reasonable to expect that the action of such type of inhibitors will be not affected greatly by the virus’s RBD mutations.

As we shown by FEP calculations the reduction of the S1-RBD binding to STE90-C11 was well pronounced. Thus, we also studied the structural changes due to the N501Y mutation based on most represented structure from the FEP MD ensemble. Two equivalent antibodies can bind to the virus’s spike S1-RBD. Thus, the N501Y mutation can affect the binding by both via direct interactions with only one of them and also to produce a change in the interactions between the individual STE90-C11 units.

One of the obvious alters detected was the disruption of the formed by Gln498 H-bond with Ser30. This is also valid for Thr500 – Ser30 hydrogen bond and in general the hydrophobic interactions in this part of the S1 RBD binding surface (see **Figure 3**). Indeed, the Asn501-Ser30 and Gly502-Gly28 H-bonds were also removed. The Tyr501 did not provided any significant interactions with the antibody. The Tyr58 located in the second chain of the antibody dramatically changed its conformation leaving the central point of the binding to the antibody without any stable hydrophobic stabilization and also disrupting the hydrogen bond network formed by Ser56 of the antibody. These changes were introduced because of the Tyr501 stabilization role on the conformation of Arg403 and Arg408. Tyr58-Thr415 and Ser56-Asp420 H-bonds were also altered by the conformation of Tyr58. All of these conformational changes were not observed during the wild type simulation. These data should be further confirmed by long term MD simulations which are underway in our lab and more details would be revealed.

**Figure 3.**
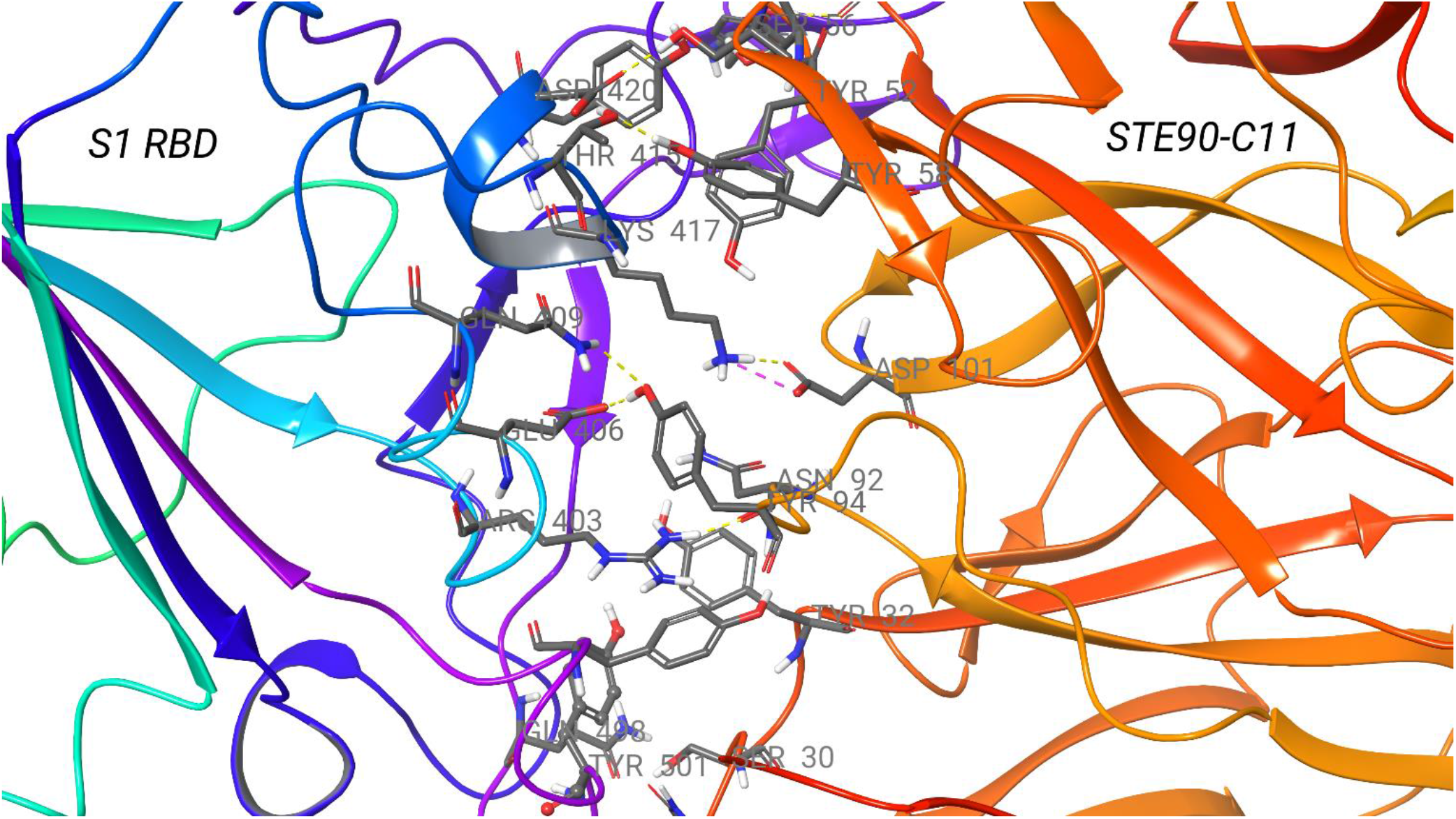
The identified changes in the interactions between S1 RBD and STE90-C11 human derived antibody due to the N501Y mutation. With yellow dot lines are shown the H-bonds.

The mechanism of action of K417N mutation is also clear (see **Figure 3**). After the disruption of the aftermentioned interactions of S1 RBD with STE90-C11, and in particular the conformational change of Tyr58, the Lys417 occupied the same area making a strong hydrogen bond with Asp101. The cancellation of this H-Bond by the asparagine mutation and the other established interactions of Lys417 are an obvious reason for the further decrease in the binding. More MD simulations would be helpful to reveal the additive mechanism of action of N501Y and K417N mutations. The same is valid also for the description of the S1 RBD – ACE2 interactions. Based on the 500 ns-long MD simulation of N501Y mutant we can conclude the Lys417 seems have a compensatory mechanism of action increasing the free energy by 1.6 kcal/mol, as shown per our FEP calculations. It has an important role in the binding and can create a strong H-bonds with Asp30 and His34. However, this is not the real case when both N501Y and K417N mutations are present at the same time and their effect on the S1 RBD conformational changes and S1 RBD-ACE2 binding remains to be revealed.

## Acknowledgments

Thanks due to the Suman Sirimulla for the helpful discussion and Nikolay Tsvetanov from Micar21 and Apolo LLC for the financial support. We thank also to Dimitar Dimitrov for the management of several activities.

## References

[1] K. Kupferschmidt. Mutant coronavirus in the United Kingdom sets off alarms, but its importance remains unclear. Science, December 20, 2020; https://www.sciencemag.org/news/2020/12/mutant-coronavirus-united-kingdom-sets-alarms-its-importance-remains-unclear

[2] H. Tegally, E. Wilkinson, M. Giovanetti, A. Iranzadeh & et.al. Emergence and rapid spread of a new severe acute respiratory syndrome-related coronavirus 2 (SARS-CoV-2) lineage with multiple spike mutations in South Africa. medRxiv 2020, 2020.12.21.20248640; doi: https://doi.org/10.1101/2020.12.21.20248640

[3] Gu H, Chen Q, Yang G, et al. (2020) Adaptation of SARS-CoV-2 in BALB/c mice for testing vaccine efficacy. Science 2020, 369, 1603–1607.

[4] Kemp S, Datir R, Collier D, et al. Recurrent emergence and transmission of a SARS-CoV-2 Spike deletion ΔH69/V70. bioRxiv. 2020:2020.2012.2014.422555.

[5] Rambaut A, Loman N, Pybus O, et al. Preliminary genomic characterisation of an emergent SARS-CoV-2 lineage in the UK defined by a novel set of spike mutations. https://virological.org/t/preliminary-genomic-characterisation-of-an-emergent-sars-cov-2-lineage-in-the-uk-defined-by-a-novel-set-of-spike-mutations/563. Published 2020. Accessed 22 December, 2020.

[6] Wang, L.; Wu, Y.; Deng, Y; Kim, B.; Pierce, L.; Krilov, G.; Lupyan, D.; Robinson, S.; Dahlgren, M.K.; Greenwood, J.; et al. Accurate and Reliable Prediction of Relative Ligand Binding Potency in Prospective Drug Discovery by Way of a Modern Free-Energy Calculation Protocol and Force Field J. Am. Chem. Soc., 2015, 137 (7), 2695–2703.

[7] Abel, R.; Wang, L.; Harder, E.D.; Berne, B.J.; Friesner, R.A., “Advancing Drug Discovery through Enhanced Free Energy Calculations,” Acc. Chem. Res., 2017 50 (7), 1625–1632.

[8] A. J. Clark, C. Negron, K. Hauser, M. Sun, L. Wang, R. Abel, R. A. Friesner. Relative Binding Affinity Prediction of Charge-Changing Sequence Mutations with FEP in Protein–Protein Interfaces, J. Mol. Biol., 2019, 431, 7, 1481–1493.

[9] Clark AJ, Gindin T, Zhang B, Wang L, Abel R, Murret CS, Xu F, Bao A, Lu NJ, Zhou T, Kwong PD, Shapiro L, Honig B, Friesner RA. Free Energy Perturbation Calculation of Relative Binding Free Energy between Broadly Neutralizing Antibodies and the gp120 Glycoprotein of HIV-1. J Mol Biol. 2017; 429(7):930–947.

[10] Ford MC, Babaoglu K. Examining the Feasibility of Using Free Energy Perturbation (FEP+) in Predicting Protein Stability. J. Chem. Inf. Model. 2017 57(6):1276–1285.

[11] Duan J, Lupyan D, Wang L. Improving the Accuracy of Protein Thermostability Predictions for Single Point Mutations. Biophys J. 2020119(1):115–127.

[12] Fratev F, Sirimulla S. An Improved Free Energy Perturbation FEP+ Sampling Protocol for Flexible Ligand-Binding Domains. Sci. Rep. 2019; 9(1):16829.

[13] Harder, E.; Damm, W.; Maple, J.; Wu, C.; Reboul, M.; Xiang, J.Y.; Wang, L.; Lupyan, D.; Dahlgren, M.K.; Knight, J.L.; Kaus, J.W.; Cerutti, D.S.; Krilov, G.; Jorgensen, W.L.; Abel, R.; Friesner, R.A., OPLS3: A Force Field Providing Broad Coverage of Drug-like Small Molecules and Proteins. J. Chem. Theory Comput, 2016, 2 (1), 281–296

[14] F. Bertoglio, V. Fühner, M. Ruschig, P. A. Heine, A SARS-CoV-2 neutralizing antibody selected from COVID-19 patients by phage display is binding to the ACE2-RBD interface and is tolerant to known RBD mutations. BioRxiv, 2020, December 3, doi: https://doi.org/10.1101/2020.12.03.409318

[15] Fratev F, Miranda-Arango M, Lopez AB, Padilla E, Sirimulla S. Discovery of GlyT2 Inhibitors Using Structure-Based Pharmacophore Screening and Selectivity Studies by FEP+ Calculations. ACS Med. Chem. Lett. 2019; 10(6):904–910.

[16] Fratev F, Steinbrecher T, Jónsdóttir SÓ. Prediction of Accurate Binding Modes Using Combination of Classical and Accelerated Molecular Dynamics and Free-Energy Perturbation Calculations: An Application to Toxicity Studies. ACS Omega. 2018; 3(4):4357–4371.

[17] Starr, T. N., Greaney, A. J., Hilton, S. K., Crawford, K., Navarro, M. J., Bowen, J. E., Tortorici, M. A., Walls, A. C., Veesler, D., & Bloom, J. D. (2020). Deep mutational scanning of SARS-CoV-2 receptor binding domain reveals constraints on folding and ACE2 binding. Cell 2020, 182, 5, 3, 1295–1310.e20

[18] Cantuti-Castelvetri L, Ojha R, Pedro LD, Djannatian M, Franz J, Kuivanen S, van der Meer F, Kallio K, Kaya T, Anastasina M, Smura T, Levanov L, Szirovicza L, Tobi A, Kallio-Kokko H, Österlund P, Joensuu M, Meunier FA, Butcher SJ, Winkler MS, Mollenhauer B, Helenius A, Gokce O, Teesalu T, Hepojoki J, Vapalahti O, Stadelmann C, Balistreri G, Simons M. Neuropilin-1 facilitates SARS-CoV-2 cell entry and infectivity. Science. 2020; 370(6518):856–860.

